# Functional analyses of *STIM1* mutations reveal a common pathomechanism for tubular aggregate myopathy and Stormorken syndrome

**DOI:** 10.1101/663088

**Authors:** Georges Arielle Peche, Coralie Spiegelhalter, Roberto Silva-Rojas, Jocelyn Laporte, Johann Böhm

## Abstract

Tubular aggregate myopathy (TAM) is a progressive disorder essentially involving muscle weakness, cramps, and myalgia. TAM clinically overlaps with Stormorken syndrome (STRMK), associating TAM with miosis, thrombocytopenia, hyposplenism, ichthyosis, short stature, and dyslexia. TAM and Stormorken syndrome arise from gain-of-function mutations in *STIM1* or *ORAI1*, both encoding key regulators of Ca^2+^ homeostasis, and mutations in either gene results in excessive Ca^2+^ entry. The pathomechanistic similarities and differences of TAM and Stormorken syndrome are only partially understood. Here we provide functional *in cellulo* experiments demonstrating that STIM1 harboring the TAM D84G or the STRMK R304W mutation similarly cluster and exert a dominant effect on the wild-type protein. Both mutants recruit ORAI1 to the clusters, induce major nuclear import of the Ca^2+^-dependent transcription factor NFAT, and trigger the formation of circular membrane stacks. In conclusion, the analyzed TAM and STRMK mutations have a comparable impact on STIM1 protein function and downstream effects of excessive Ca^2+^ entry, highlighting that TAM and Stormorken syndrome involve a common pathomechanism.

## INTRODUCTION

Tubular aggregate myopathy (TAM) and Stormorken syndrome (STRMK) are spectra of the same disease affecting muscle, platelets, spleen, and skin (1, 2). The majority of the TAM patients primarily present with muscle weakness, cramps, and myalgia with a heterogeneous age of onset and disease severity (3–6). Additional features including thrombocytopenia, hyposplenism, miosis, ichthyosis, short stature, hypocalcemia, or dyslexia can be seen as well, and the occurrence of the totality of the multi-systemic signs constitutes the diagnosis of Stormorken syndrome (7–18).

Both TAM and Stormorken syndrome are characterized by the presence of densely packed membrane tubules in muscle fibers, and investigations on muscle sections by immunofluorescence have shown that these tubular aggregates contain various sarcoplasmic reticulum (SR) proteins, suggesting that they originate from the SR (4, 7, 15, 16, 19). The tubular aggregates are the histopathological hallmark of TAM and Stormorken syndrome, and were also described in hypokalemic periodic paralysis, myasthenic syndrome, malignant hyperthermia, inflammatory or ethyltoxic myopathy, and accumulate in normal muscle with age (20–25).

TAM and Stormorken syndrome are caused by heterozygous missense mutations in *STIM1* (4, 10–12) (OMIM #605921) or *ORAI1* (12) (OMIM #610277), both encoding key factors of store-operated Ca^2+^ entry (SOCE). SOCE is a major mechanism regulating Ca^2+^ homeostasis and thereby drives a multitude of Ca^2+^-dependent cellular functions including muscle contraction. STIM1 has a single transmembrane domain and is primarily localized at the endoplasmic/sarcoplasmic reticulum with an N-terminal luminal part containing the Ca^2+^-sensing EF hands, and a C-terminal cytosolic part containing coiled-coil domains (CC1-3). Upon Ca^2+^ store depletion, STIM1 undergoes a conformational change, clusters in vicinity to the plasma membrane, and recruits and activates the Ca^2+^ channel ORAI1 to trigger extracellular Ca^2+^ entry and reticular Ca^2+^ store refill (26–28). Functional experiments have shown that the *STIM1* and *ORAI1* mutations induce excessive extracellular Ca^2+^ entry despite replete reticular Ca^2+^ stores (4, 10–12, 15).

Fourteen different *STIM1* mutations have been described in patients with TAM and Stormorken syndrome, including 12 mutations in the luminal EF-hands, and two mutations in the cytosolic CC1 domain (3–14, 18). Patients with EF-hand mutations mainly manifest a muscle phenotype and only isolated multi-systemic features, while the most common R304W substitution in the cytosolic CC1 domain, found in thirteen unrelated families, was essentially described with the full clinical picture of Stormorken syndrome (7, 10–14).

The shared genetic causes of TAM and Stormorken syndrome, the consistent skeletal muscle histopathology, and the overlapping clinical presentation of affected individuals raises the possibility of a common sequence of events leading to either TAM or STRMK. In an attempt to elucidate and compare the cellular defects underlying both disorders, we performed a series of functional and comparative *in cellulo* experiments. We demonstrate that both the TAM D84G and the STRMK R304W mutation similarly impact on STIM1 migration and clustering at the plasma membrane, the interaction with ORAI1, the nuclear translocation of the Ca^2+^-dependent transcription factor NFAT, and the formation of membrane stacks. We therefore conclude that TAM and Stormorken syndrome involve a common pathomechanism.

## MATERIALS AND METHODS

### Constructs

The human YFP-STIM1, mCherry-STIM1, ORAI1-eGFP, and eGFP-NFAT constructs were kind gifts from Nicolas Demaurex (University of Geneva, Switzerland), Richard S. Lewis (Stanford University, USA), Liangyi Chen (Beijing University, China), and Cristina Ulivieri, (Universita degli studi di Siena, Italy), respectively. The STIM1 (c.251A>G; p.D84G, c.910C>T; p.R304W) and ORAI1 (c.319G>A; p.V107M) point mutations were introduced by site-directed mutagenesis using the Pfu DNA polymerase (Stratagene, La Jolla, USA).

### Cell culture

C2C12 murine myoblasts were cultured in DMEM, supplemented with 20% fetal calf serum (FCS) and 0.5% gentamycin (all Gibco Life Technologies, Carlsbad, USA), grown at 37°C and 5% CO_2_, and transfected using Lipofectamine® 2000 (Invitrogen Life Technologies Carlsbad, USA) at 50% confluency in Opti-MEM (Gibco Life Technologies). HeLa cells were grown in DMEM with 5% FCS and 0.5% gentamycin and transfected at 70% confluency using Lipofectamine® 2000. Co-expression experiments were conducted using a 1:1 plasmid ratio.

Twenty-four hours post transfection, cells seeded on glass slides were fixed using 4% paraformaldehyde (PFA) for 20 min at RT, treated with 50mM ammonium chloride (NH4Cl) for 15 min and rinsed in 1xPBS. Nuclei were stained with DAPI (Sigma Aldrich), and the coverslips were mounted using FluorSave reagent (Calbiochem, Darmstadt, Germany). Cells were classified according to the cytosolic or nuclear localization of the eGFP-NFAT signal, and the statistical significance was assessed through one-way ANOVA followed by Dunnett’s post-hoc test. All experiments were performed in triplicate and monitored with the Leica TCS SP8 AOBS inverted confocal microscope equipped with a Leica HCX PL APO CS2 63x/1.4 oil immersion objective (Leica, Wetzlar, Germany).

### Total Internal Reflection Fluorescence (TIRF) microscropy

The TIRF plane of acquisition was determined according to the YFP fluorescence detected at the plasma membrane, and the images were acquired at resting state and following addition of 2μM thapsigargin (Sigma-Aldrich, Saint Louis, USA) using a Nikon TI-eclipse inverted microscope equipped with a 100x, 1.49 NA oil-immersion objective (Nikon, Tokyo, Japan).

### Correlative Light and Electron Microscopy (CLEM)

Cells were cultured on micro-patterned aclar supports (29), transfected with WT or mutant YFP-STIM1. Cells were precisely located and imaged by confocal microscopy (Leica TCS SP2-AOBS), and then chemically fixed with 0.1M sodium cacodylate buffer containing 2.5% paraformaldehyde and 2.5% glutaraldehyde, and post-fixed in 1% osmium tetroxide for 1h at 4°C. After extensive washing in distilled water, cells were incubated in 1% uranyl acetate for 1h at 4°C, dehydrated through a graded series of ethanol solutions, and embedded in epoxy resin polymerized 48h at 60°C. Ultrathin sections (60 nm) were mounted on pioloform-coated slot grids and examined with a Philips CM12 (80 kV) electron microscope equipped with a Gatan ORIUS 1000 CCD camera (FEI Company Hillsboro, USA).

## RESULTS

### TAM and STRMK mutants constitutively cluster at the plasma membrane

We first assessed the effect of the TAM and STRMK-related *STIM1* mutations on the SOCE-dependent accumulation of STIM1 in vicinity to the plasma membrane. To this aim, we transfected C2C12 myoblasts with WT or mutant (TAM D84G or STRMK R304W) YFP-STIM1, and monitored the fluorescence by TIRF microscopy. Cells expressing WT STIM1 showed a diffuse signal inside the cell at the resting state, and addition of 2 μM thapsigargin induced Ca^2+^-store depletion and the formation of STIM1 clusters at the plasma membrane (Figure 1a). This is in agreement with previous studies showing that STIM1 migrates to the plasma membrane and clusters upon SOCE activation (30). In contrast, cells transfected with STIM1 D84G or R304W displayed clusters at the plasma membrane independently of thapsigargin treatment. This supports the idea that both TAM and STRMK-related *STIM1* mutations involve a gain-of-function resulting in constitutive STIM1 clustering despite replete Ca^2+^ stores.

**Figure 1:**
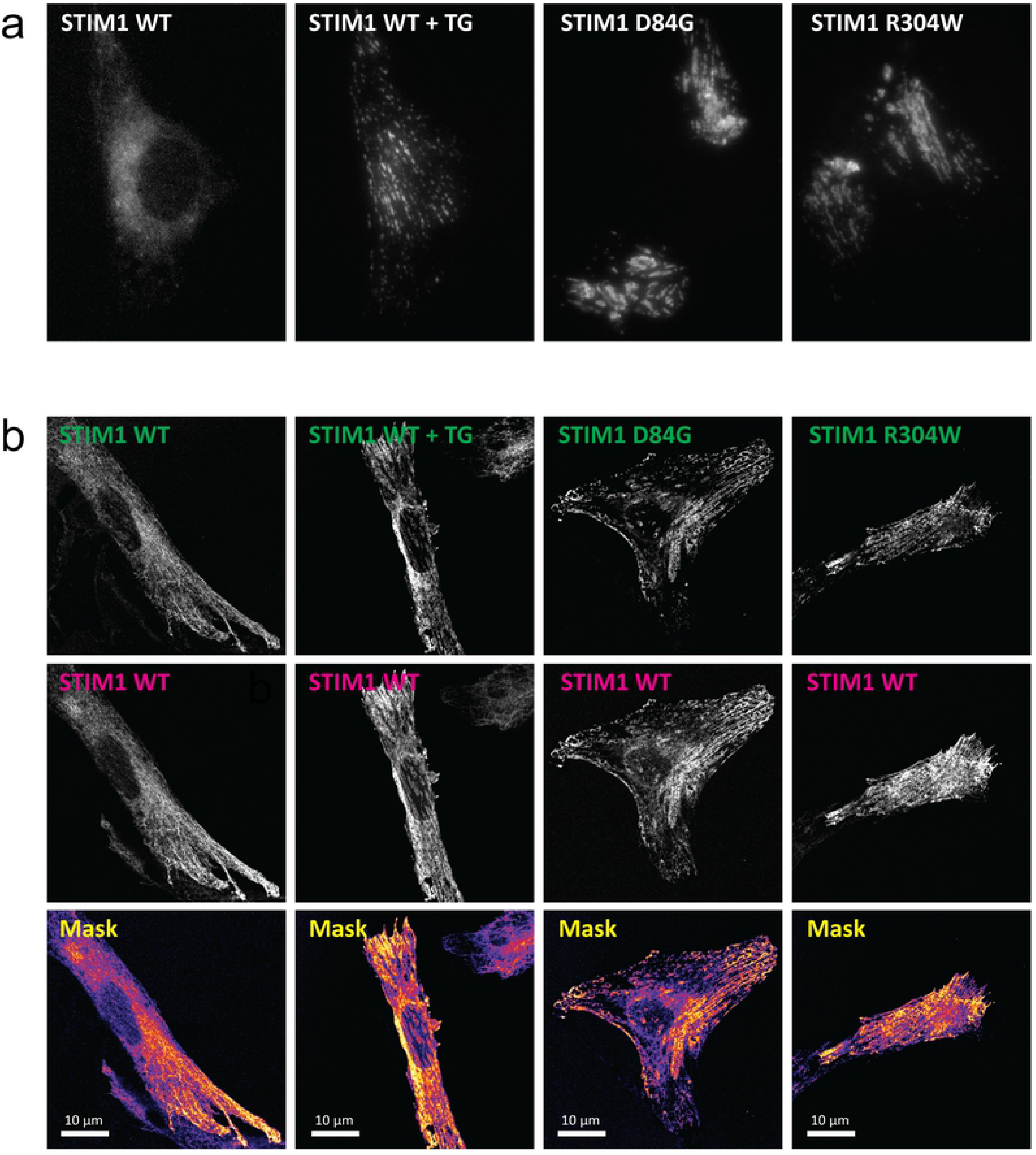
Impact of TAM and STRMK mutations on STIM1 clustering. (a) TIRF microscopy on transfected C2C12 muscle cells demonstrates that wild type STIM1 has a diffuse localization inside the cell, and clusters at the vicinity of the plasma membrane upon Ca^2+^ store depletion through thapsigargin (TG). In contrast, both TAM D84G and STRMK R304W mutants cluster without TG treatment and independently of the reticular Ca^2+^ load. (b) Co-transfection of C2C12 cells with differentially tagged STIM1 constructs shows that D84G and R304W STIM1 sequester wild type STIM1 to the clusters, revealing a dominant effect.

### TAM and STRMK mutants sequester WT STIM1

Both TAM and Stormorken syndrome result from heterozygous *STIM1* missense mutations (4, 10–12). To investigate a potential effect of mutant STIM1 on the wild type protein, we co-transfected C2C12 cells with WT or mutant YFP-STIM1 and WT mCherry-STIM1. Cells co-expressing differently tagged WT proteins displayed diffuse overlapping signals. Addition of thapsigargin provoked co-clustering of both WT proteins, illustrating that the fluorescent YPF and mCherry tags do not impact on the intracellular localization or clustering of STIM1 (Figure 1b). Cells co-expressing D84G or R304W with WT STIM1 exhibited major STIM1 clusters of overlapping YFP and mCherry signals without thapsigargin treatment. These results demonstrate that both TAM and STRMK mutants recruit wild type STIM1 to the clusters without Ca^2+^ store depletion, and thereby evidence a dominant effect leading to the constitutive clustering of WT STIM1.

### TAM and STRMK mutants recruit ORAI1 and are less sensitive to Ca^2+^ influx

The cytosolic coiled-coil domains of STIM1 contain the STIM1-ORAI1 activating domain (SOAR), which is essential for the interaction with the plasma membrane Ca^2+^ channel ORAI1 (26, 31, 32). To examine the effect of the TAM and STRMK mutations on the recruitment of ORAI1 to the STIM1 clusters, we transfected C2C12 cells with WT or mutant mCherry-STIM1 and WT ORAI1-eGFP (Figure 2). When co-expressed, WT STIM1 and WT ORAI1 significantly co-cluster, and we also observed clusters containing STIM1 and ORAI1 in cells expressing mutant D84G or R304W STIM1 and WT ORAI1. To investigate the influence of Ca^2+^ entry on the recruitment of ORAI1 to the STIM1 clusters, we next co-transfected HeLa cells with WT or mutant mCherry-STIM and mutant V107M ORAI1-eGFP (Figure 2). The V107M mutation affects an essential amino acid of the pore-forming unit of ORAI1, and has been shown to generate a channel with constant Ca^2+^ permeability (15, 33). Major clusters were found in more than 80% of the cells expressing mutant STIM1 and mutant ORAI1, demonstrating that the leaky V107M ORAI1 channel is largely recruited to the D84G or R304W STIM1 clusters. In contrast, less than 10% of the cells expressing wild type STIM1 and mutant ORAI1 displayed prominent clusters. These data illustrate that steady Ca^2+^ inflow through V107M ORAI1 efficiently disassembles or prevents the formation of wild type STIM1 clusters and thereby inactivates SOCE, while the mutation-induced STIM1 clusters persist independently of the Ca^2+^ flux through the CRAC channel.

**Figure 2:**
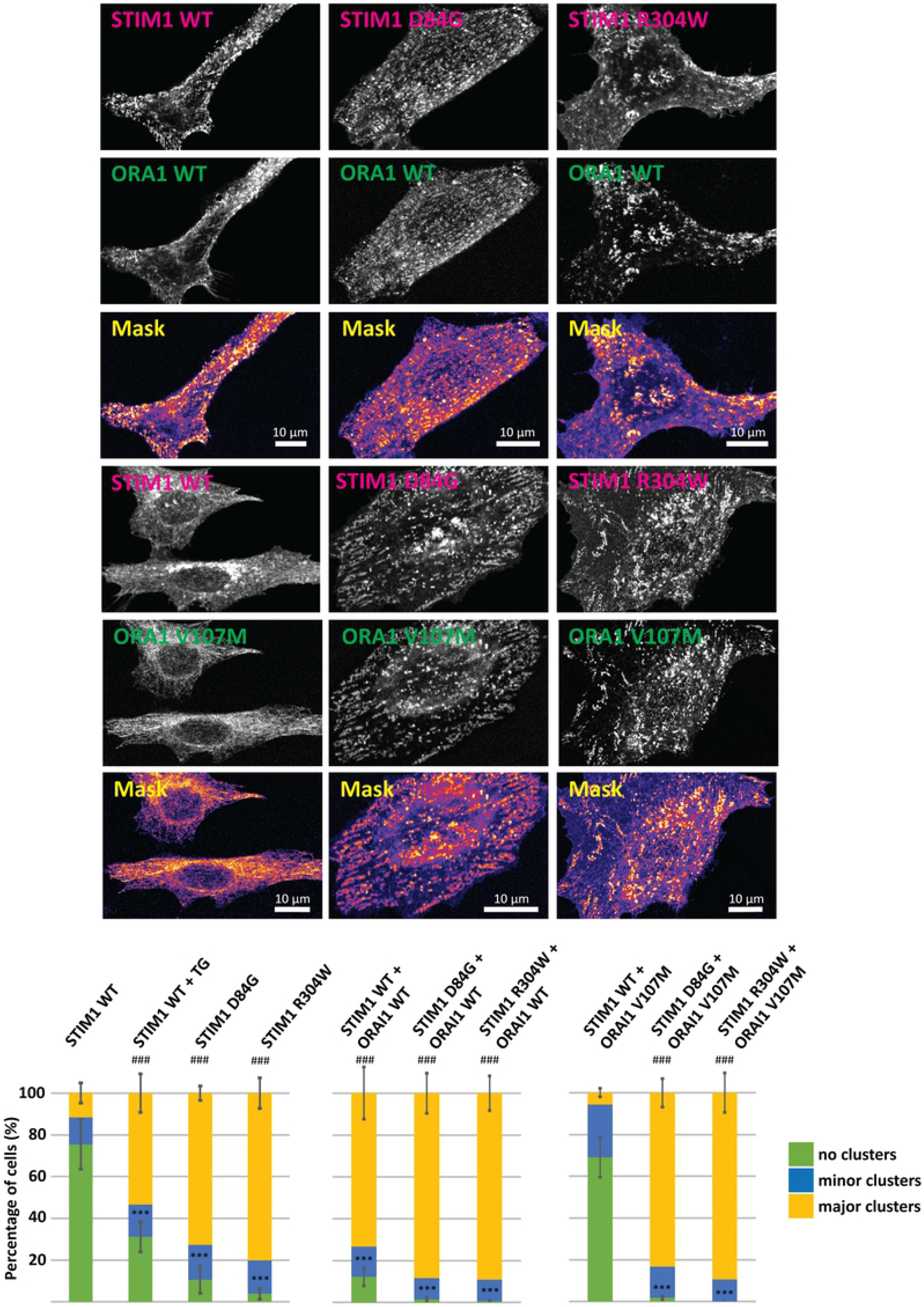
Impact of the TAM and STRMK mutations on ORAI1 recruitment. Confocal microscopy on co-transfected C2C12 cells shows that wild-type STIM1 co-localizes with wild-type ORAI1 in the clusters, but no clusters are seen in cells expressing wild-type STIM1 and the leaky V107M ORAI1 channel. In contrast, both STIM1 mutants D84G and R304W co-localize with V107M ORAI1 in clusters. For quantification, between 100 and 200 cells were counted per experiment and per condition. The total number of counted cells was set to 100%, the percentage of cells with major STIM1/ORAI1 clusters corresponds to the yellow bar, cells with minor cluster to the blue bar, and cells without clusters to the green bar. Error bars represent the standard deviation (SD), and the statistical difference compared to the first column of the set is indicated by * for cells without clusters and by # for cells with major clusters. P-value < 0.001:*** or ###.

### TAM and STRMK mutants increase nuclear NFAT translocation

NFATc2 (nuclear factor of activated T cells) belongs to the NFAT family of transcription factors. It has a cytosolic localization when inactive, and gets dephosphorylated through calcineurin upon increased cytosolic Ca^2+^ levels, resulting in nuclear import (34). The ratio of nuclear versus cytoplasmic NFAT therefore reflects the Ca^2+^ concentration in the cytosol and represents a suitable parameter to assess the downstream effect of excessive Ca^2+^ entry resulting from *STIM1* mutations. In accordance, previous studies have shown that constitutively active STIM1 induces major NFAT translocation to the nucleus (35). To assess if the *STIM1* TAM and STRMK mutations similarly promote nuclear NFAT import, we co-transfected WT or mutant mCherry-STIM1 with eGFP-NFAT and quantified the intracellular NFAT localization (Figure 3). About 21% of the cells expressing WT STIM1 showed a nuclear NFAT localization. Following caffeine treatment, known to induce Ca^2+^ release from the reticulum to the cytosol, nuclear NFAT was observed in 73% of the cells. Cells exogenously expressing D84G or R304W STIM1 displayed major nuclear NFAT translocation in the absence of caffeine treatment, with intense nuclear GFP signals in 58% and 61% of the cells, respectively. Quantification of the NFAT signal in the cytoplasm versus the nucleus revealed a ratio of 1.8 for cells expressing WT STIM, a ratio of 0.4 when treated with caffeine, and a ratio of 0.1 for cells expressing mutant D84G or R304W STIM1, illustrating that NFAT is mostly nuclear in cells transfected with mutant STIM1. Taken together, the STIM1 mutations D84G and R304W similarly and significantly promote nuclear NFAT import, demonstrating a downstream effect of cellular Ca^2+^ excess induced by *STIM1* mutations.

**Figure 3:**
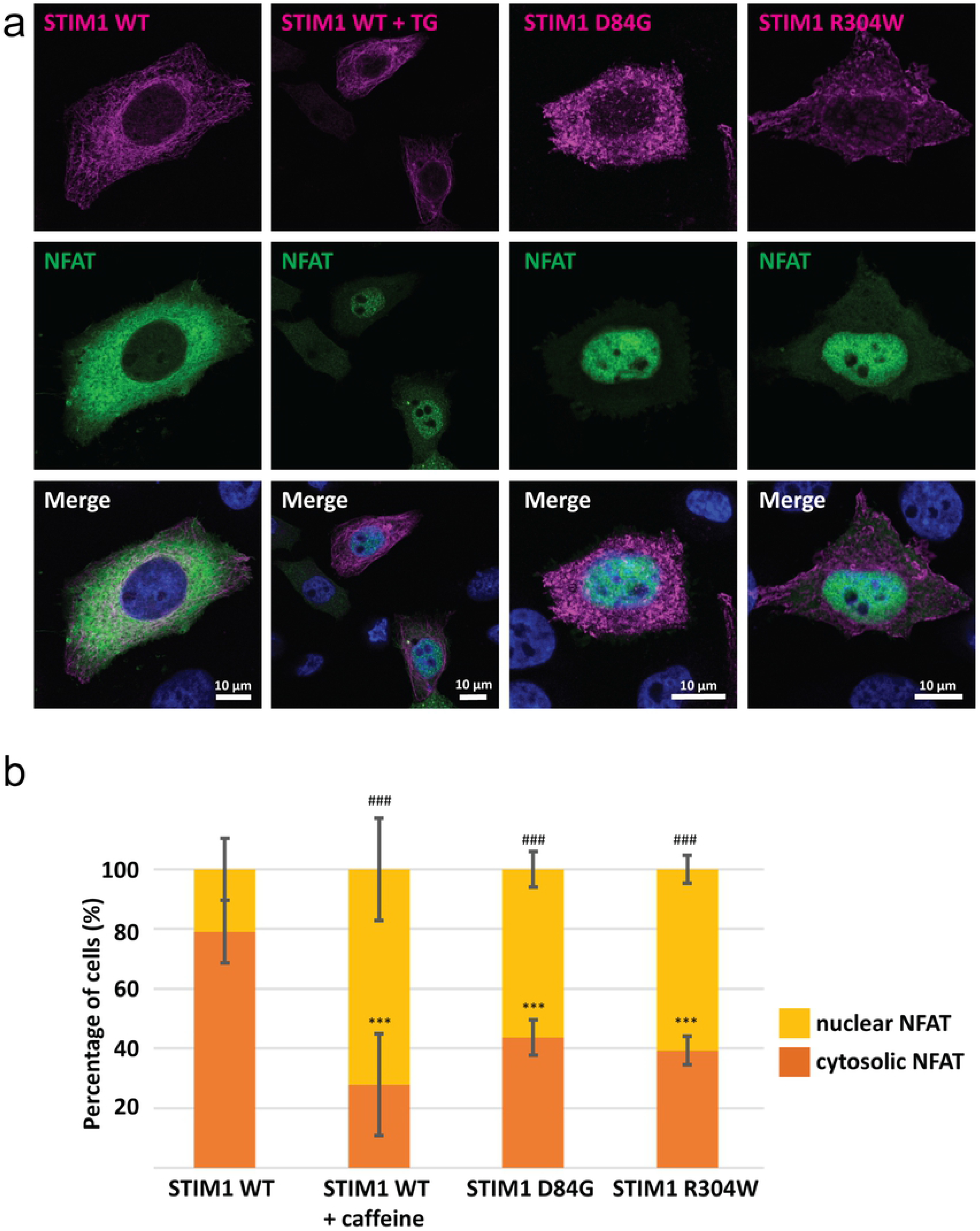
Impact of the TAM and STRMK mutations on NFAT localization. Confocal microscopy on transfected HeLa cells and quantification shows a nuclear NFAT signal in 21% and 73% of cells expressing WT STIM1 before and after caffeine treatment. In untreated cells expressing D84G or R304W STIM1, nuclear NFAT was seen in 58% and 61% of the cells, respectively. For quantification, between 100 and 200 cells per condition were counted in three independent experiments. The total number of counted cells was set to 100%, and the bars show the percentage of cells with nuclear NFAT in yellow and the percentage of cells with cytosolic NFAT in orange. Error bars represent the standard deviation. The statistical differences compared to cells transfected with WT STIM1 are indicated by * for nuclear NFAT and # for cytoplasmic NFAT. P-value < 0.001:*** or ###.

### TAM and STRMK mutants promote circular membrane stack formation

Tubular aggregates are regular arrays of single- or double-walled membrane tubules appearing as honeycomb-like structures on transverse muscle sections, and are the histopathological hallmark of TAM and Stormoken syndrome (1, 2, 20, 36). To investigate the tubular aggregate formation *in cellulo*, we performed correlated light and electron microcopy (CLEM) on HeLa cells transfected with WT or mutant YFP-STIM1 constructs. We observed membrane stacks in cells exogenously expressing D84G or R304W STIM1, but not in cells transfected with wild-type STIM1 (Figure 4). These membrane stacks were most often found with a circular shape and connected with the endoplasmic reticulum. Our observations illustrate that STIM1 harboring amino acid substitutions found in TAM and Stormorken syndrome patients induce the emergence of aberrant membrane accumulations, potentially representing the first step of tubular aggregate formation.

**Figure 4:**
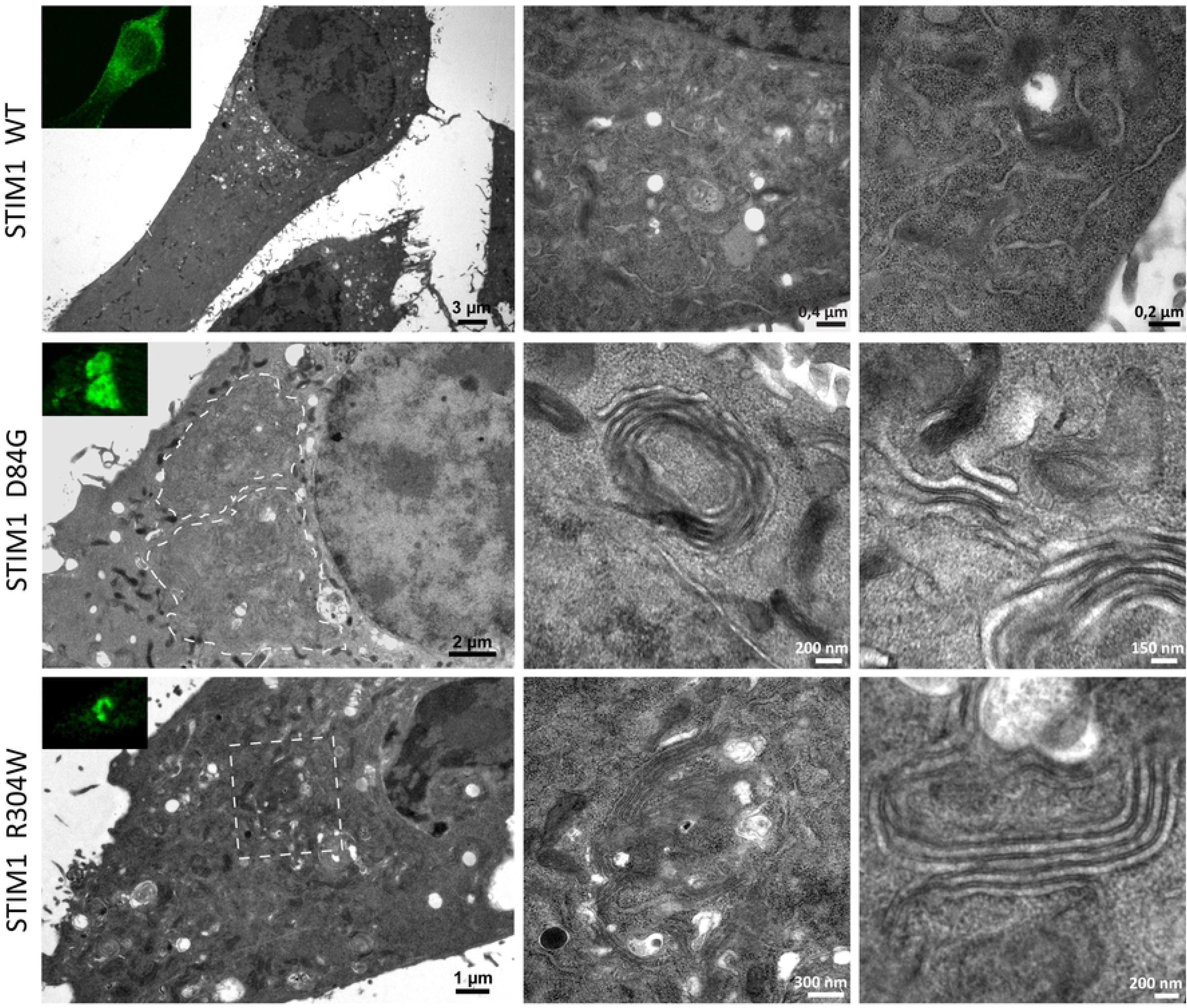
Impact of the TAM and STRMK mutations on membrane architecture. CLEM allows the analysis of the same cell by light and electron microscopy. In areas corresponding to fluorescent STIM1 cluster, we observed circular membrane stacks in HeLa cells exogenously expressing the STIM1 mutants D84G or R304W, but not in cells transfected with WT STIM1.

## DISCUSSION

In this study, we investigated the molecular impact of *STIM1* mutations associated with tubular aggregate myopathy (TAM) and Stormorken syndrome (STRMK) at different levels of the SOCE pathway. We and others have previously shown that the TAM and STRMK mutations induce constitutive STIM1 clustering and activation of the Ca^2+^ entry channels (4, 10–12). Here we addressed and compared the sequence of events leading to the cellular defects of TAM and Stormorken syndrome, and we found that the TAM D84G and STRMK R304W mutants similarly impact on STIM1 clustering, ORAI1 recruitment, Ca^2+^-dependent nuclear translocation of NFAT, and the formation of circular membrane stacks.

### The STIM1 mutations have multiple effects on cluster formation

Through TIRF experiments we showed that both TAM D84G and STRMK R304W STIM1 mutants constitutively cluster in the vicinity of the plasma membrane and recruit ORAI1 to the clusters although the amino acid substitutions affect different STIM1 protein domains. The EF-hand D84G mutation is believed to directly or indirectly impair Ca^2+^ sensing, resulting in STIM1 unfolding, clustering, and exposure of the SOAR domain mediating the interaction with ORAI1 (1, 3). The luminal R304W mutation does not interfere with Ca^2+^ binding. Instead, a recent study revealed that it induces a helical elongation within the coiled-coil domain, and thereby promotes STIM1 clustering and the exposure of the SOAR domain (37). In both cases, the mutations induce constitutive ORAI1 binding and activation, resulting in major extracellular Ca^2+^ influx despite replete Ca^2+^ stores (4, 10–12).

In agreement with previous studies (26), we observed that exogenously expressed wild-type STIM1 and ORAI1 co-localize and cluster in cells. However, co-expression of wild-type STIM1 and V107M ORAI1, previously described to generate a leaky channel with reduced Ca^2+^ selectivity (15, 33), leads to a diffuse reticular distribution of STIM1. We therefore conclude that the constant ion entry through the permeable ORAI1 V107M channel either dissipates the STIM1 clusters or prevents their formation. It is known that massive Ca^2+^ entry generates hyperpolarized potentials at the intracellular channel gate, and thereby promotes a rapid inactivation of ORAI1 and the dissociation of STIM1 from ORAI1 (Ca^2+^-dependent inactivation, CDI)(38, 39). In contrast to wild-type STIM1, both TAM and STRMK mutants efficiently recruited V107M ORAI1 to the clusters. This suggests that the interaction between the STIM1 mutants and the leaky ORAI1 channel is insensitive to elevated local Ca^2+^ levels, and that the consequential reduction of CDI significantly contributes to excessive extracellular Ca^2+^ entry. Consistently, decreased CDI was measured in lymphoblasts from a Stormorken syndrome patient, resulting in elevated basal Ca^2+^ levels (12).

We furthermore found that the clusters in cells co-expressing wild-type and mutant STIM1 are formed by both proteins despite replete Ca^2+^ stores. This indicates that the TAM or STRMK mutants are able to oligomerize with wild type STIM1 and to sequester the non-mutated protein to the clusters. Such an effect has not yet been described for *STIM1* mutations, and demonstrates that wild-type STIM1 contributes to the constitutive activation of ORAI1 and the excessive extracellular Ca^2+^ entry characterizing TAM and Stormorken syndrome.

### The downstream effects of excessive Ca^2+^ entry

As a consequence of constitutive STIM1 and ORAI1 activation, myoblasts from TAM/STRMK patients were found to exhibit increased basal Ca^2+^ levels in the cytosol (4, 5, 7, 10–12). Ca^2+^ is a physiological key factor triggering numerous signalling cascades including the NFAT pathway. The NFAT transcription factors mainly reside in the cytosol at low Ca^2+^ concentrations, and become phosphorylated upon rising Ca^2+^ levels and translocate to the nuclei (34). In agreement, we found that cells expressing mutant STIM1 and manifesting prominent STIM1/ORAI1 clusters also displayed major nuclear NFAT translocation as compared to cells expressing wild-type STIM1. This is consistent with previous reports showing that the exogenous expression of constitutively active STIM1 causes nuclear NFAT import (35), while silencing of STIM1 prevents NFAT translocation (40). In this context it is interesting to note that *STIM1* belongs to the NFAT target genes (41), suggesting a positive feedback controlling SOCE that might modulate the disease development of TAM and Stormorken syndrome.

Tubular aggregates are the histopathological hallmark in skeletal muscle of patients with tubular aggregate myopathy and Stormorken syndrome (20, 42, 43). Although it is not fully understood how the tubular aggregates form, they were shown to contain sarcoplasmic reticulum proteins and large amounts of Ca^2+^ (19), and are therefore likely to be the consequence of excessive Ca^2+^ storage in the reticulum. Accordingly, constitutive SOCE activation was shown to induce the formation of membrane stacks in transfected cells (44), and our CLEM experiments on cells expressing mutant STIM1 identified multilayer reticulum membranes of circular shape, which may represent the first step of tubular aggregate formation. Of note, tubular aggregates were not seen in the TAM/STRMK mouse model harbouring the STIM1 R304W mutation and clinically recapitulating the human disorder (45), demonstrating that tubular aggregate formation is species-specific and rather a consequence than a cause of the disease.

### Common pathomechanisms in tubular aggregate myopathy and Stormorken syndrome

Tubular aggregate myopathy essentially affects skeletal muscle, and a subset of TAM patients harboring STIM1 EF-hand mutations additionally manifested one or several signs of Stormorken syndrome (3–5, 8, 9, 18). Conversely, Stormorken syndrome patients carrying the most common R304W mutation usually present the full clinical picture of TAM, miosis, thrombocytopenia, hyposplenism, ichthyosis, short stature, and dyslexia (7, 10–14), but individual patients with muscle weakness as the main clinical sign were also reported (9). This illustrates that TAM and Stormorken syndrome form a clinical continuum, and raises the question on the underlying common and diverging pathomechanisms. Here we investigated and compared the SOCE pathway defects in cells expressing STIM1 mutants, and we uncovered that the TAM and STRMK mutants had a comparable effect on cluster formation, ORAI1 recruitment, nuclear NFAT translocation, and membrane rearrangements, indicating that TAM and Stormorken syndrome involve a common pathomechanism. The phenotypic differences between both disorders might result from a different mutational impact on fast Ca^2+^-dependent inactivation (CDI), corresponding to the inactivation of ORAI1 through high local Ca^2+^ levels. Indeed, electrophysiological studies have shown that the R304W mutant suppresses fast CDI, while STIM1 harboring a luminal amino acid substitution was indistinguishable from the wild-type (12). This suggests that R304W, but not the EF-hand mutations causes a prolonged Ca^2+^ influx and might account for the significant aberrations in multiple tissues in patients with Stormorken syndrome. Alternatively, especially the cytosolic STIM1 R304W mutation might have an additional pathogenic effect on other interaction partners as the non-selective TRPC channels (46), and the full clinical picture of Stormorken syndrome would then result from the excessive entry of Ca^2+^ and other cations in the different tissues.

## Concluding remarks

Here we demonstrate that missense mutations in different STIM1 domains have a similar pathogenic effect on SOCE and downstream pathways. Our finding that *STIM1* mutations exert a dominant effect points to a suitable therapeutic approach for TAM and Stormorken syndrome through an allele-specific downregulation of *STIM1* expression. The resulting reduction of the STIM1 protein level is thereby not expected to be pathogenic as heterozygous carriers of *STIM1* loss-of-function mutations, associated with immunodeficiency, are healthy(2).

## ACKNOWLEDGEMENTS

We thank Catherine Koch and Pascal Kessler for their outstanding technical help, and Nicolas Demaurex (University of Geneva, Switzerland), Richard S. Lewis (Stanford University, USA), Liangyi Chen (Beijing University, China), and Cristina Ulivieri, (Universita degli studi di Siena, Italy) for providing the plasmids.

## AUTHOR CONTRIBUTIONS

JL and JB designed the study and supervised the research, GAP and CS performed the experiments, GAP, CS, RSR, and JB analyzed the data, GAP, JL, and JB wrote the manuscript.

